# Preparation of Asymmetric Vesicles with Trapped CsCl Avoids Osmotic Imbalance, Non-Physiological External Solutions, and Minimizes Leakage

**DOI:** 10.1101/2021.05.03.442494

**Authors:** Ming-Hao Li, Daniel P. Raleigh, Erwin London

## Abstract

The natural asymmetry of cellular membranes influences their properties. In recent years, methodologies for preparing asymmetric vesicles have been developed that rely on the methyl-α-cyclodextrin catalyzed exchange of lipids between donor lipid multilamellar vesicles and acceptor lipid unilamellar vesicles, and the subsequent separation of the, now asymmetric, acceptor vesicles from the donors. Isolation is accomplished by pre-loading acceptor vesicles with a high concentration of sucrose, typically 25% (w/w), and separating from donor and cyclodextrin by sucrose gradient centrifugation. We found that when the asymmetric vesicles were dispersed under hypotonic conditions using physiological salt solutions, there was enhanced leakage of an entrapped probe, 6-carboxyfluorescein. Studies with symmetric vesicles showed this was due to osmotic pressure and was specific to hypotonic solutions. Inclusion of cholesterol partly reduced leakage but did not completely eliminate it. To avoid having to use hypotonic conditions or to suspend vesicles at non-physiological solute concentrations to minimize leakage, a method for preparing asymmetric vesicles using acceptor vesicle-entrapped CsCl at a physiological salt concentration (100 mM) was developed. Asymmetric vesicles prepared with the entrapped CsCl protocol were highly resistant to 6-carboxyfluorescein.

## INTRODUCTION

Natural biological membranes are inherently asymmetric, and their asymmetry influences their properties ^1^. In recent years, an efficient methodology for preparing asymmetric vesicles has been developed which opens the door to the biophysical studies of the effects of asymmetry on membrane properties in a controlled environment ^2–8^. The method involves the cyclodextrin-catalyzed exchange of lipids between donor multilamellar vesicles and acceptor unilamellar vesicles, and the subsequent separation of the, now altered, acceptor vesicles from the donors. In many cases, isolation is accomplished by pre-loading the acceptor vesicles with sucrose and separating by sucrose gradient centrifugation ^2–7^. To achieve this, a significant concentration of sucrose, often 25% (w/w) (all sucrose concentrations are given in % (w/w)), is encapsulated in the interior of the asymmetric vesicles. This, in turn, leads to a significant difference in osmolarity between the interior and exterior of the vesicle if the vesicles are suspended in a physiological salt solution, such as phosphate-buffered saline (PBS). We found this osmotic imbalance enhances leakage of 6-carboxyfluorescein from vesicles, which interferes with studies of protein-induced leakage. Inclusion of cholesterol in vesicles partly, but not fully, suppressed this leakage.

In principle, issues due to the osmotic imbalance could be overcome by including 25% (w/w) sucrose in the bulk buffer on the exterior of the vesicles. However, via crowding, or by effects on solution viscosity, high sucrose concentrations can influence the properties of proteins including their stability, folding rates, as well as their tendency to aggregation and to form amyloid ^9–15^. A method in which donor vesicles, rather than acceptor vesicles are entrapped with sucrose and then pelleted to separate them from acceptor vesicles, followed by filtration to remove cyclodextrin can avoid these problems ^8^. However, that method would be difficult to use when acceptor membranes have a high density due to the use of lipids that promote tighter lipid packing, such as cholesterol, or due to inclusion of proteins in the acceptor vesicles.

Here we demonstrate and validate a simple and robust alternative approach using vesicle entrapped CsCl. This allows the preparation of asymmetric vesicles with no osmotic pressure imbalance dispersed in physiological salt. We found vesicles prepared this way were highly resistant to leakage of 6-carboxyfluorescein.

## EXPERIMENTAL SECTION

### Materials

Cholesterol, 1-palmitoyl-2-oleoyl-sn-glycero-3-phospho-L-serine (sodium salt) (POPS), 1-palmitoyl-2-oleoyl-sn-glycero-3-phosphoethanolamine (POPE), 1-myristoyl-2-{6-[(7-nitro-2-1,3-benzoxadiazol-4-yl)amino]hexanoyl}-sn-glycero-3-phosphocholine (NBD-PC), and 1-palmitoyl-2-oleoyl-glycero-3-phosphocholine (POPC) were purchased from Avanti Polar Lipids (Alabaster, AL). Lipids were dissolved in chloroform and stored at −20°C. The concentrations of unlabeled lipids were determined by dry weight, and the NBD-PC concentration was determined by using absorbance with ε_NBD-PC_ 22,000 M^−1^ cm^−1^ at 464 nm. 5(6)-Carboxyfluorescein was purchased from Sigma-Aldrich. Methyl-α-cyclodextrin (MαCD) was purchased from Arachem (Tilburg, the Netherlands). MαCD was dissolved in 20 mM tris 100 mM NaCl pH 7.4 buffer at concentration close to 300 mM, and then filtered through a Sarstedt (Nümbrecht, Germany) 0.2 μm pore syringe filter. The concentration of MαCD was determined by comparing the refractive index of MαCD solutions to MαCD solutions with known concentrations. 1-(4-Trimethylammoniumphenyl)-6-phenyl-1,3,5-hexatriene p-toluenesulfonate (TMADPH) was purchased from the Molecular Probes division of Invitrogen. TMADPH was dissolved in ethanol. The TMADPH concentration was determined by using absorbance with εTMADPH 74,100 M^−1^ cm^−1^ at 365 nm. Thin layer chromatography plates (Silica Gel 60) were purchased from VWR International (Batavia, IL).

### Preparation of symmetric large unilamellar vesicles (LUVs)

Lipids initially dissolved in chloroform were mixed. The lipid mixture was dried by a stream of nitrogen and subject to high vacuum for one hour. Multilamellar vesicles (MLV) were prepared by dissolving the dry lipid mixture in buffer (20 mM tris 100 mM NaCl at pH 7.4) and incubating in a 55°C shaker for 15 min. LUVs were prepared from MLVs with 7 freeze-thaw cycles followed by 11 extrusions with a 100 nm poly-carbonate membrane.

### Membrane permeability assays

The same method was used to prepare LUVs for membrane permeability assays, except that the dry lipid film was dissolved in solutions containing 80 mM carboxyfluorescein. Free carboxyfluorescein was removed using a PD-10 desalting column. 400 μM lipid solutions of LUVs were loaded into a Hellma 96 well quartz plate. Fluorescence of carboxyfluorescein was measured using a DTX 880 plate reader at 25°C (excitation wavelength: 485 nm, and emission wavelength: 535 nm). The fluorescence change was calculated as following: Normalized fluorescence intensity = (F(t)-F(0))/(Fmax-F(0)). F(0) is fluorescence at time zero, F(t) is the fluorescence intensity of carboxyfluorescein at a given time t, and Fmax is the fluorescence intensity after disrupting LUV, achieved by the addition of 2% (v/v) triton x-100.

### Preparation of asymmetric LUVs using MαCD

To prepare asymmetric LUVs with POPE (30%): POPS (30%): cholesterol (40%) in the inner leaflet and POPC (60%): cholesterol (40%) in the outer leaflet, (all lipid concentrations are mol% unless otherwise noted), 8 mM acceptor LUV (POPE (30%): POPS (30%): cholesterol (40%)) was prepared as above with 20 mM tris 100 mM CsCl, or 20 mM tris 100 mM NaCl 25% (w/w) sucrose at pH 7.4. 16 mM donor MLV (100% POPC) was prepared using 20 mM tris 100 mM NaCl buffer with 40 mM MαCD. 1ml acceptor LUV was diluted with 10 mM tris 100 mM NaCl and centrifuged to remove CsCl or sucrose (188,965 g, 37,500 rpm, 45 min, Beckman L8-80M, SW-60 rotor). The acceptor LUV pellet was resuspended in 1ml 20 mM tris 100 mM NaCl pH 7.4 buffer. Donor MLV was incubated in a 55°C shaker for 2 hours and then mixed with resuspended acceptor LUV. The 1ml acceptor and 1ml donor mixture was incubated in a 37°C shaker for 45 min. CsCl-containing acceptor/donor mixture was loaded on top of a 7% (w/w) sucrose solution and sucrose-containing acceptor/donor mixture was loaded on top of a 9% (w/w) sucrose solution. Acceptor and donor were separated by using centrifugation (188,965 g, 37,500 rpm, 45 min, Beckman L8-80M, SW-60 rotor). After centrifugation, the pellet was resuspended in 10 mM tris and 20 mM NaCl pH 7.4 buffer. Lipid concentration was determined by using thin layer chromatography (TLC).

### Thin Layer Chromatography to monitor lipid composition

TLC plates were preheated at 100°C and cooled to room temperature prior to use. Standard lipid mixtures (different amounts of cholesterol:POPE:POPC:POPS) and asymmetric membrane samples were loaded on the TLC plate. 65:25:5 chloroform:methanol: ammonia hydroxide (v/v) was used to separate each lipid. After air-drying for 10 min, the plate was sprayed with a 3% (w/v) cupric acetate, 8% (v/v) phosphoric acid solution, dried for 45 min, and charred at 180°C for 2–5 min. The charred plate was scanned using an Epson 1640XL scanner (Epson America Inc., Long Beach, CA) and analyzed by ImageJ. Standard curves were generated for each lipid by plotting the intensity of the TLC bands vs lipid weight and were fitted using linear regression in Microsoft Excel. Lipid composition ratios and concentrations of asymmetric LUVs were calculated using the standard curves.

### Determination of POPS in the outer leaflet of LUV using TMADPH assays

50 μM lipid LUVs were mixed with 0.25 μM of TMADPH in 20 mM tris 100 mM NaCl pH 7.4 buffer to measure fluorescence (F) (excitation: 358 nm; emission: 427 nm). 0.25 μM of TMADPH was dissolved in 100% ethanol to normalize to fluorescence in organic solvent (F_0_) (excitation: 358 nm; emission: 427 nm). Fluorescence intensity were recorded in a 1 cm pathlength cell on a Horiba QuantaMaster Spectrofluorimeter (Horiba Scientific, Edison, NJ). Symmetric LUVs (POPC (60%): cholesterol (40%); POPS (5%): POPE (5%): POPC (50%): cholesterol (40%); POPS (10%): POPE (10%): POPC (40%): cholesterol (40%); POPS (15%): POPE (15%): POPC (30%): cholesterol (40%); POPS (20%): POPE (20%): POPC (20%): cholesterol (40%); and POPS (30%): POPE (30%): cholesterol (40%)) were used to generate a standard curve of F/Fo vs mole % POPS. F/Fo of asymmetric LUVs was compared to the standard curve.

## RESULTS and DISCUSSION

### The effect of an osmotic pressure imbalance on membrane permeability

To examine the effect of osmotic pressure on membrane permeability, a set of carboxyfluorescein dye leakage assays were performed with three different conditions: approximately isotonic, hypertonic, and hypotonic (figure 1) using symmetric vesicles comprised of pure POPC. The approximately isotonic is actually modestly hypotonic because the interior of the vesicles contains 80 mM carboxyfluorescein. However, we refer to these conditions as “near isotonic” in the rest of this report. Under near isotonic conditions (figure 1), low to moderate leakage was observed over 40 hours for both 100 mM NaCl inside and outside, and 25% (w/w) sucrose inside and outside (figure 1A, black circles, figure 1B and 1C, gray triangles). In contrast, under hypotonic conditions with 25% (w/w) sucrose inside and 100 mM NaCl outside relatively high leakage was observed (figure 1A, gray triangles). Under hypertonic conditions (figure 1B and figure 1C, black circles) low to moderate leakage was observed. This implies that vesicles may have better ability to tolerate squeezing than stretching. Notice that similar behavior was observed for the hypertonic conditions when vesicles were suspended in either 25% (w/w) sucrose or 500 mM NaCl. Also, as illustrated in figure 1B and 1C, there was no difference between near isotonic and hypertonic samples even though the former contained a high sucrose concentration inside. This demonstrates that sucrose inside the vesicles is not responsible for a high degree of leakage and that the effects are instead due to the osmotic imbalance.

**Figure 1.**
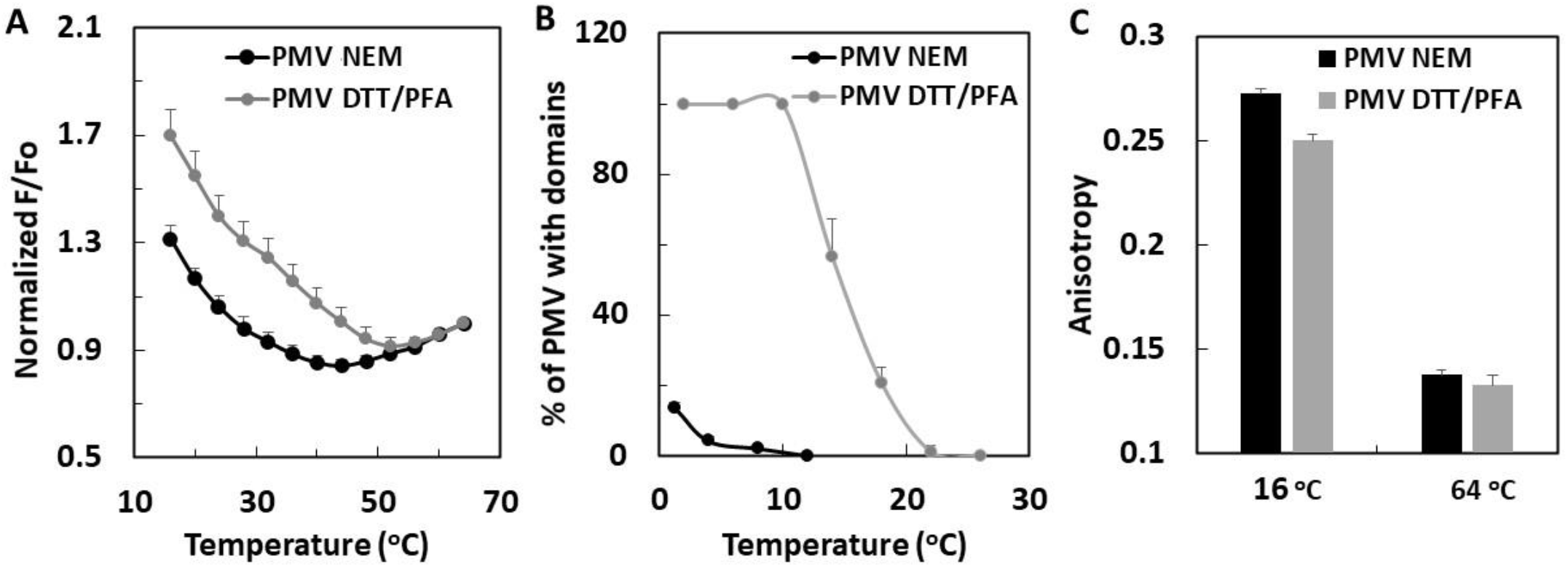
Effect of an osmotic pressure imbalance across the lipid bilayer and the effect of sucrose on membrane permeability. Carboxyfluorescein fluorescence intensity using vesicles with (open circles) and without (closed symbols) 2% (v/v) triton x-100 are shown. Various buffers were used outside and inside the vesicles. Outside: (A) tris 20 mM, NaCl 100 mM, (B) tris 20 mM, NaCl 100 mM, 25% (w/w) sucrose, (C) tris 20 mM, NaCl 500 mM; Inside: (black circles), tris 20 mM, NaCl 100 mM; (gray triangles), tris 20 mM, NaCl 100 mM, 25% (w/w) sucrose. Lipid composition was 100% POPC and the lipid concentration was 400 μM (all lipid concentrations are mol% unless otherwise noted). The concentration of carboxyfluorescein was 80 mM inside the vesicle. The error bars are smaller than the symbols. In this and all the following experiments, unless otherwise noted average (mean) values and standard deviations from three samples are shown.

### Cholesterol reduces the effect of an osmotic pressure imbalance on membrane permeability

Cholesterol is an essential component of the cell membrane and can modulate membrane permeability induced by membrane active peptides or other perturbants ^16–19^. Consequently, we examined the impact of osmotic pressure on the permeability of model LUV’s containing cholesterol. Membrane permeability of 100% POPC vesicles was significantly higher vesicles comprised of POPC (60%): cholesterol (40%) (figure 2), which implies that the LUV’s without cholesterol are leakier in the presence of osmotic pressure than those with cholesterol.

**Figure 2.**
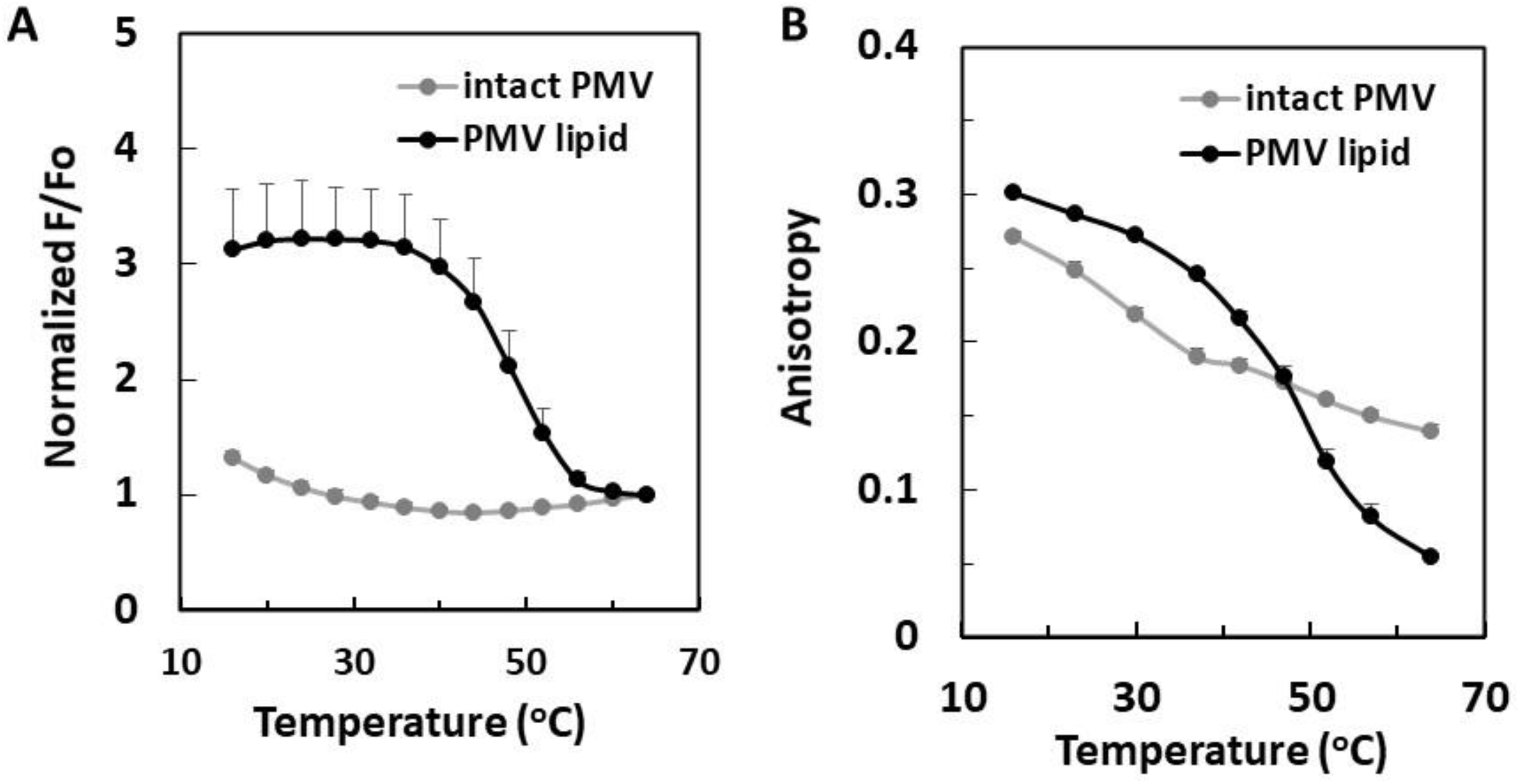
Cholesterol reduces the permeability of symmetric membranes under osmotic pressure. Carboxyfluorescein fluorescence intensity using vesicles with (open circles) and without (closed symbols) 2% (v/v) triton x-100 are shown. Symmetric LUV’s compositions were (A) 100% POPC or (B) POPC (60%): cholesterol (40%). 80 mM carboxyfluorescein and 25% (w/w) sucrose was encapsulated in all samples. Experiments were conducted in tris 20 mM NaCl 100 mM pH 7.4 buffer at 25°C.

### Preparation of asymmetric vesicles without an osmotic pressure imbalance

Next, 100 mM CsCl was used to replace 25% sucrose entrapped inside lipid vesicles. Because these molecules are entrapped in vesicles to allow their purification of acceptor vesicles from donor vesicles after a lipid exchange experiment, we first optimized the method for acceptor vesicle purification. Acceptor vesicles with entrapped CsCl and donor MLV were loaded on top solutions with various sucrose concentrations (3~9%), separately. Next, the acceptor LUV and the donor MLV were pelleted using high-speed centrifugation (189,000g, 45 min, table 1). Table 1 shows centrifugation at low external sucrose concentrations resulted in a high level of pelleting of donor vesicles, while centrifugation at high external sucrose concentrations resulted in a low yield of acceptor vesicles. Using 7% sucrose, acceptor vesicles were pelleted under conditions which give a low level of donor vesicle contamination (purity >95%) and a reasonable yield (~10%), which is similar to the sucrose encapsulation method ^4^.

**Table 1.** Recoveries of separate donor and acceptor samples using different sucrose cushions (3, 6, 7, 8, or 9% (w/w) of sucrose) and high-speed centrifugation (188,965 g, 37,500 rpm, 45 min). Acceptor was loaded with 20 mM tris 100 mM CsCl and donor was loaded with 20 mM tris 100 mM NaCl. The initial concentrations of donor and acceptor are 16 mM and 8 mM, respectively. Acceptor purities are calculated using the formula: [acceptor]/([donor+acceptor])*100%. Lipid composition for acceptor LUVs and donor MLVs are POPS (29.75%): POPE (75%): cholesterol (40%): NBD-PC (0.5%) and POPC (99.5%): NBD-PC (0.5%), respectively.

| Sucrose | 3% | 6% | 7% | 8% | 9% |
| --- | --- | --- | --- | --- | --- |
| **Acceptor Yield %** | 65.4 ± 19.5 | 25.6 ± 13.5 | 9.94 ± 1.57 | 8.89 ± 3.39 | 4.87 ± 2.39 |
| **Donor Yield %** | 8.03 ± 10.1 | 1.97 ± 1.26 | 0.20 ± 0.23 | 0.05 ± 0.03 | 0.12 ± 0.08 |
| **Purity (%)**<br>**acceptor/(acceptor+donor)** | 83.0 ± 19.3 | 86.3 ± 7.08 | 96.4 ± 4.40 | 98.8 ± 0.29 | 93.1 ± 7.64 |

The effect of varying the type of salt (NaCl or CsCl) entrapped upon vesicle yield was also examined. If the effect of CsCl is due to its high density, we expect it should give higher vesicle yields. Table 2 shows that, except for POPC vesicles, loading vesicles with CsCl gave higher recoveries after centrifugation over sucrose (189,000g, 45 min, and 6 ~ 7% of sucrose) compared to vesicles loaded with NaCl. However, POPC vesicles were not sufficiently dense to pellet even when loaded with CsCl. This shows that the identity of the lipid used is a variable that can also affect yield.

**Table 2.** Effect of salt density and lipid composition on the % pelleted for different LUVs after centrifugation in a sucrose cushion. LUVs were loaded with 20 mM tris 100 mM NaCl or 20 mM tris 100 mM CsCl and pelleted with high-speed centrifugation (188,965 g, 37,500 rpm, 45 min). Initial concentrations of LUVs were 8 mM.

| Lipid composition | Salt inside vesicles | % Pelleted After Centrifugation in 6 to 7% (w/w) Sucrose |  |
| --- | --- | --- | --- |
|  |  | 6% | 7% |
| POPS (50%): POPE (50%) | CsCl | 57.3 ± 2.4 | 41.7 ± 3.5 |
|  | NaCl | 5.3 ± 0.6 | 5.4 ± 0.6 |
| POPS (30%): POPC (30%): Cholesterol (40%) | CsCl | 25.6 ± 13.5 | 9.9 ± 1.5 |
|  | NaCl | 3.6 ± 0.9 | 1.2 ± 0.2 |
| 100% POPC | CsCl | 0.9 ± 0.3 | 0.1 ± 0.04 |
|  | NaCl | 2.0 ± 1.3 | 0.2 ± 0.2 |

### Carboxyfluorescein leakage for LUVs prepared with CsCl depends upon osmotic pressure

Symmetric vesicles with the similar osmolarity in the interior and exterior did not exhibit a high level of carboxyfluorescein leakage when using NaCl or CsCl (figure 3). This was true even though CsCl was used at a much higher concentration than NaCl. This shows that CsCl by itself does not promote carboxyfluorescein leakage. However, considerable leakage was observed with hypotonic conditions (800 mM CsCl inside and 100 mM NaCl outside) (figure 3). Thus, in agreement with the data in figure 1, it appears that hypotonic conditions promote carboxyfluorescein leakage.

**Figure 3.**
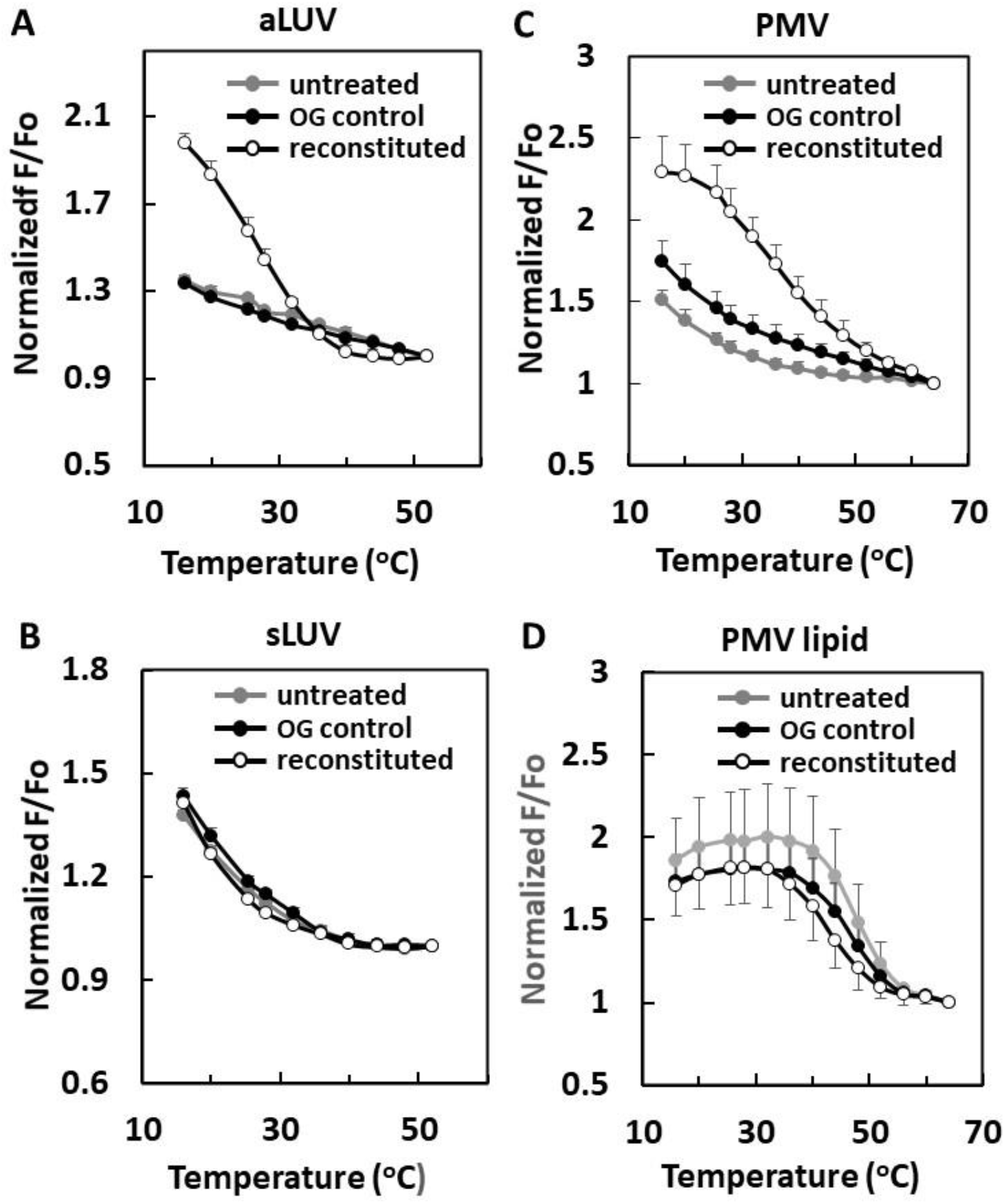
Effect of an osmotic pressure imbalance on membrane permeability for 100% POPC vesicles prepared using the CsCl protocol. Carboxyfluorescein fluorescence intensity using vesicles with (open circles) and without 2% (v/v) triton x-100 (closed symbols) are shown. Various buffers were used inside or outside the vesicle. (black circles), inside: tris 20 mM, NaCl 100 mM, outside: tris 20 mM, NaCl 100 mM; (gray squares), inside: tris 20 mM, CsCl 800 mM, outside: tris 20 mM, NaCl 800 mM; (gray triangle), inside: tris 20 mM CsCl 800 mM, outside: tris 20 mM, NaCl 100 mM. The total lipid concentration for each LUV was 400 μM. The concentration of carboxyfluorescein was 80 mM inside the vesicle.

### Carboxyfluorescein leakage is low for asymmetric LUVs prepared using CsCl

We next examined carboxyfluorescein leakage in asymmetric LUVs prepared using entrapped CsCl under near-isotonic conditions. Asymmetric vesicles were prepared with a 1:1 ratio of POPE:POPS in their inner leaflet and POPC in the outer leaflet, with or without 40 mol% cholesterol (which locates in both leaflets ^20^). In both the presence and absence of cholesterol, asymmetric LUVs prepared using near isotonic conditions showed low carboxyfluorescein membrane permeability (figure 4).

**Figure 4.**
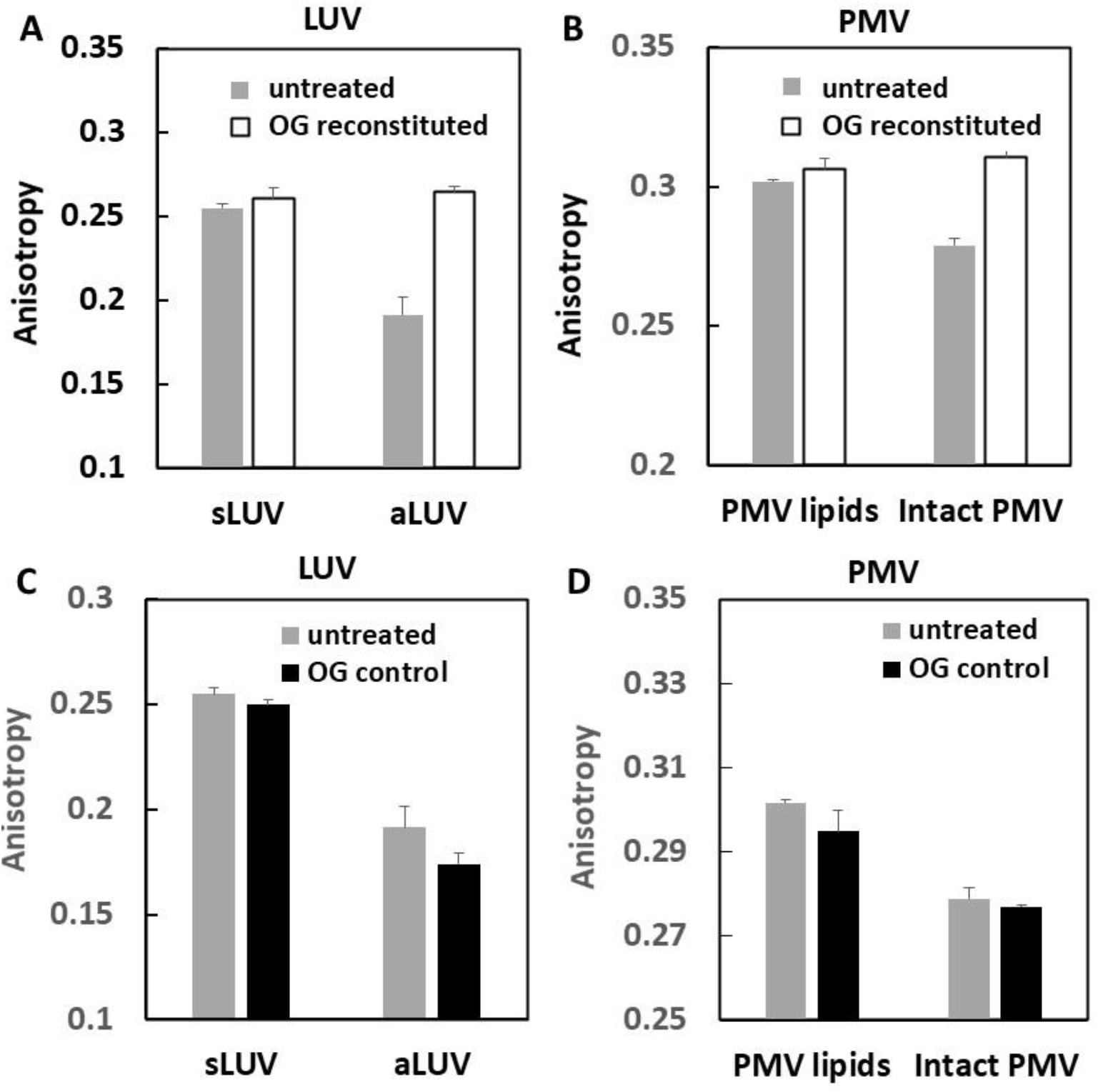
Very low carboxyfluorescein leakage is observed from asymmetric vesicles prepared using the CsCl protocol. Carboxyfluorescein fluorescence intensity using vesicles with (open circles) and without (black circles) 2% (v/v) triton x-100 are shown. The total lipid concentration was 400 μM. Asymmetric LUVs were loaded with 20 mM tris 100 mM CsCl and 80 mM carboxyfluorescein. The experiments were conducted in tris 20 mM, NaCl 100 mM, pH 7.4 buffer at 25°C. (A) Asymmetric vesicles prepared from POPS (50%): POPE (50%) acceptor and POPC (100%) donor. (B) Asymmetric vesicles prepared from POPS (30%): POPE (30%): cholesterol (40%) donor and POPC (100%) acceptor.

It should be noted that the use of CsCl did not perturb the ability to prepare vesicles that were asymmetric or to obtain efficient exchange. We used the TMADPH fluorescence assay to monitor LUV asymmetry ^4^. TMADPH is a cationic probe, and its binding to the outer leaflet of lipid vesicles is enhanced by the presence of the anionic lipid POPS in the outer leaflet ^4^. Asymmetry was assessed in samples in which the acceptor vesicles contained 1:1 POPE:POPS and 40 mol% cholesterol and the donor containing POPC. After exchange, asymmetric LUV’s should contain POPS only in the inner leaflet. In agreement with this, essentially no POPS was detected in the outer leaflet demonstrating that the asymmetry of lipid composition was achieved (figure 5). Analysis by TLC also indicated that the outer leaflet of POPS and POPE was replaced by POPC. The calculated total lipid composition from TLC was POPE (35.8 ± 2.7%): POPS (27.7 ± 2.3%): POPC (36.5 ± 1.8%) (figure S1) equivalent to 73% efficient lipid exchange in which the outer leaflet POPE:POPS is replaced by POPC, assuming about half of the phospholipid is in the outer leaflet of the LUV ^3^. (Note that MαCD does not exchange cholesterol so the exchanged acceptor vesicles also retain the original cholesterol). If higher efficiencies are desired, the ratio of donor to acceptor LUV can be increased.

**Figure 5.**
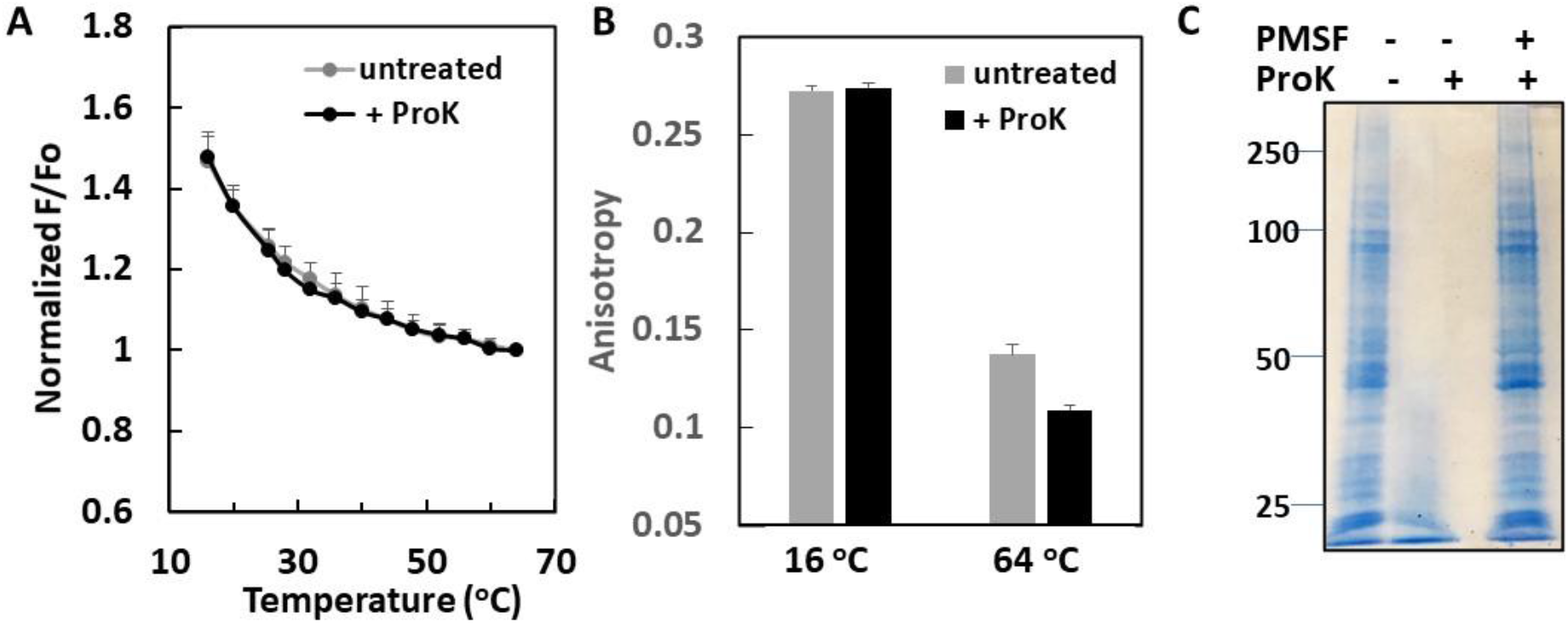
Demonstration that the CsCl protocol allows preparation of asymmetric vesicles. The POPS content of the outer leaflet of the asymmetric LUV’s was estimated by measuring the fluorescence intensity of 0.25 μM TMADPH added to the vesicles. F is the fluorescence of LUVs with TMADPH (total lipid concentration =50 μM lipid, with 0.2 μM of TMADPH), F_0_ is the fluorescence of 0.2 μM of TMADPH in ethanol. (A) Standard curve of F/F_0_ vs. mol% POPS (B) Values of F/F_0_for LUV prepared using CsCl before (gray) and after (white) MαCD exchange. The calculated mole% of POPS in the outer leaflet is −7.8 ± 7.0%. The lipid composition (mol%) before exchange was POPS (30%): POPE (30%): cholesterol (40%). The target lipid composition for the outer leaflet after exchange was POPC (60%): cholesterol (40%). Lipid compositions (mol %) for the standard curve were from left to right are POPC (60%): cholesterol (40%); POPS (5%): POPE (5%): POPC (50%): cholesterol (40%); POPS (10%): POPE (10%): POPC (40%): cholesterol (40%); POPS (15%): POPE (15%): POPC (30%): cholesterol (40%); POPS (20%): POPE (20%): POPC (20%): cholesterol (40%); and POPS (30%): POPE (30%): cholesterol (40%).

## CONCLUSIONS

Preparation of asymmetric vesicles requires conditions that give a good yield without contamination from donor lipid vesicles. Methods dependent upon density differences between the asymmetric and donor vesicles have been used to achieve this. In some cases, increasing donor lipid density, combined with filtrations is best ^8^. In other cases, increasing acceptor lipid density is necessary. When sucrose entrapped within the asymmetric vesicle is used, the vesicles must either be suspended in a hypotonic physiological solution or a non-physiological solution. Both of these alternatives are not satisfactory for some experiments. As shown here, the osmotic imbalance induced by the sucrose encapsulation method used can lead to significant leakage of entrapped molecules for hypotonic conditions. The increased leakage was due to an osmotic imbalance and is not a specific effect of sucrose. In addition, the differences between hyper and hypotonic conditions, suggests that LUVs may be better able to tolerate squeezing than stretching. The alternative to use of hypotonic conditions with asymmetric vesicles, is use of a high solute concentration in the external solution. This overcomes the leakage problem, but can perturb interactions of proteins and lipid vesicles and influence the properties of proteins and polypeptides ^9–15, 21^. Although we found that cholesterol reduces membrane permeability under an osmotic imbalance, a general method is needed to prepare asymmetric LUV with and without cholesterol. The CsCl protocol circumvents these problems, because density is increased without use of high osmolarity in the solution inside the vesicles. We found that using a 7% sucrose cushion, the acceptor can be separated from the donor with more than 95% purity and 10 % yield for the lipids studied here.

The approach demonstrated here can be adapted to vesicles which are both less dense and denser than those studied here. If the donor were denser, then one would increase the sucrose concentration used for centrifugations and replace CsCl with CsBr or CsI in the acceptor vesicles to increase their density. It should be noted that iodide is a fluorescence quencher, and is chemically unstable under some conditions, and so maybe of limited use for some experiments.

The approach illustrated here will facilitate studies of the interaction of proteins and peptides with asymmetric vesicles and will make it possible to monitor the ability of proteins and peptides to induce leakage of asymmetric LUVs under physiological salt conditions. Studies of the interaction of antimicrobial peptides, amyloidogenic polypeptides, and other membrane active polypeptides with asymmetric vesicles should also benefit from the methodology.

## Abbreviations

LUV: large unidef-lamellar vesicle
MLV: multilamellar vesicles
MαCD: methyl-α-cyclodextrin
NBD-PC: 1-myristoyl-2-{6-[(7-nitro-2-1,3-benzoxadiazol-4-yl)amino]hexanoyl}-sn-glycero-3-phosphocholine
PBS: phosphate-buffered saline
POPC: 1-palmitoyl-2-oleoyl-glycero-3-phosphocholine
POPE: 1-palmitoyl-2-oleoyl-sn-glycero-3-phosphoethanolamine
POPS: 1-palmitoyl-2-oleoyl-sn-glycero-3-phospho-L-serine (sodium salt)
TLC: thin layer chromatography
TMADPH: 1-(4-trimethylammoniumphenyl)-6-phenyl-1,3,5-hexatriene p-toluenesulfonate
v/v: volume to volume
w/w: weight to weight

## ASSOCIATED CONTENT

Supporting Information The Supporting Information is available free of charge at: 1 figure showing a TLC of asymmetric LUVs prepared using CsCl before and after MαCD exchange.

## AUTHOR CONTRIBUTIONS

Experiments were carried out by M.-H. Li. The experiments were designed and the manuscript was written through the contributions of all authors. All authors have given approval to the final version of the manuscript.

## ACKNOWLEDGEMENTS

This work was supported by NSF grants MCB-1330259 (to D.P.R.) and NIH grant GM 122493 (to E.L.).

---

**For Table of Contents Only**

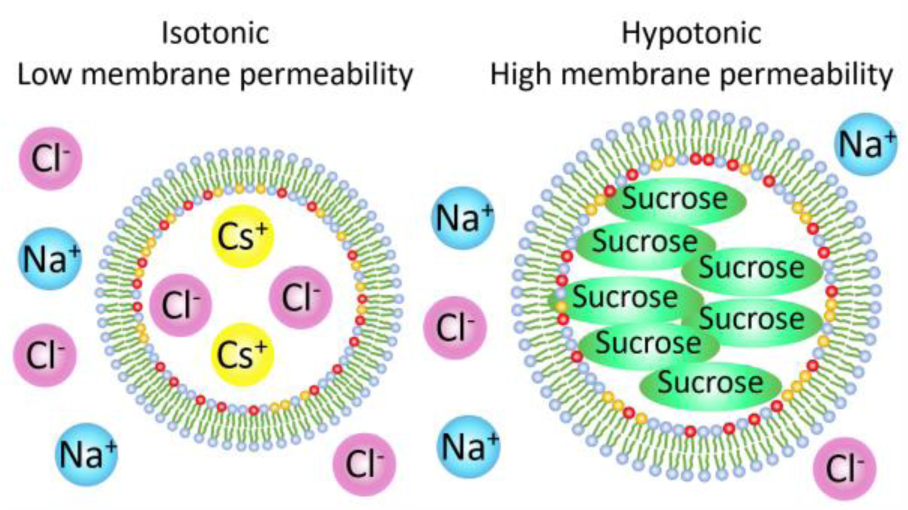

