## Supporting Information for "Preparation of Asymmetric Vesicles with Trapped CsCl Avoids Osmotic Imbalance, Non-Physiological External Solutions, and Minimizes Leakage"

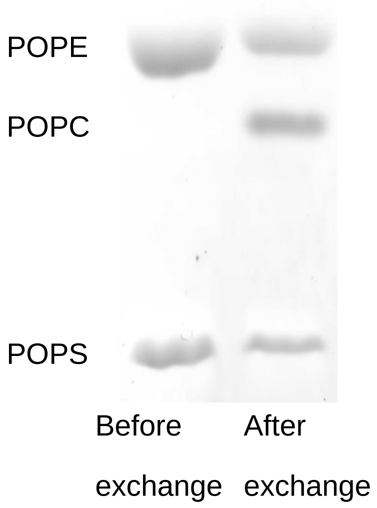


**Figure S1.** TLC analysis of asymmetric LUVs prepared using CsCl before and after MαCD exchange. Expected total phospholipid compositions of the LUVs before and after exchange were as follows. Before exchange, POPE (50%): POPS (50%); and after exchange, POPE (25%): POPS (25%): POPC (50%). Calculated phospholipid composition of the LUV before and after exchange (from three samples) was POPE (52.5 ± 11.1%): POPS (47.4 ± 11.1%) before exchange and POPE (35.8 ± 2.7%): POPS (27.7 ± 2.3%): POPC (36.5 ± 1.8%) after exchange.
